# PP2A^Rts1^ enforces a proportional relationship between cell size and growth rate

**DOI:** 10.1101/321281

**Authors:** Ricardo M. Leitao, Annie Pham, Quincy Okobi, Douglas R. Kellogg

**Affiliations:** Department of Molecular, Cell and Developmental Biology, University of California, 1156 High Street, Santa Cruz, CA 95064, USA

## Abstract

Cell size is proportional to growth rate. Thus, cells growing slowly in poor nutrients can be nearly half the size of cells growing rapidly in rich nutrients. The relationship between cell size and growth rate appears to hold across all orders of life, yet the underlying mechanisms are unknown. In budding yeast, most growth occurs during mitosis, and the proportional relationship between cell size and growth rate is therefore enforced primarily by modulating growth in mitosis. When growth is slow, the duration of mitosis is increased to allow more time for growth, yet the amount of growth required to complete mitosis is reduced, leading to birth of small daughter cells. Previous studies found that PP2A associated with the Rts1 regulatory subunit (PP2A^Rts1^) works in a TORC2-dependent feedback loop that sets cell size and growth rate to match nutrient availability. However, it was unknown whether PP2A^Rts1^ influences growth in mitosis. Here, we show that PP2A^Rts1^ is required for the proportional relationship between cell size and growth rate during mitosis. Moreover, nutrients and PP2A^Rts1^ influence the duration of mitosis, and thus the extent of growth in mitosis, via Wee1 and Pds1/securin, two conserved regulators of mitotic progression. Together, the data suggest a model in which the same global signals that set growth rate also set the critical amount of growth required for cell cycle progression, which would provide a simple mechanistic explanation for the proportional relationship between size and growth rate.

## Introduction

A critical step in the evolution of life was attainment of the capacity for growth. Along with growth, the earliest cells must have evolved mechanisms for controlling growth if they were to survive and compete. Thus, early cells needed mechanisms to control the rate and extent of such processes as membrane growth and ribosome biogenesis, while also ensuring that the rates of each process are matched to each other and to the availability of building blocks derived from nutrients. Mechanisms that control growth ultimately determine the size and shape of a cell, and are responsible for the myriad sizes, shapes, and growth rates observed in cells spanning all orders of life.

Growth of budding yeast cells illustrates how mechanisms that control cell growth define the size and shape of a cell. Growth occurs in distinct phases during the cell cycle that are characterized by different rates and patterns of growth (Johnston *et al*., 1977; McCusker *et al*., 2007; Goranov *et al*., 2009; Ferrezuelo *et al*., 2012; Leitao and Kellogg, 2017). During G1 phase, growth occurs at a slow rate and occurs over the entire surface of the cell. At the end of G1 phase, growth of the mother cell ceases and growth becomes polarized as a daughter bud emerges. Entry into mitosis triggers a 2- to 3-fold increase in the rate of growth, as well as a switch to isotropic growth that occurs over the entire bud surface. Rapid growth continues throughout mitosis and accounts for most of the volume of a yeast cell (Leitao and Kellogg, 2017). Thus, a cell growing in rich nutrient conditions achieves greater than 60% of its volume during growth in mitosis, and only 20% of its volume during G1 phase. The volume achieved during G1 phase increases to approximately 40% under poor nutrient conditions. The distinct size and shape of a budding yeast cell is ultimately defined by mechanisms that control the location and extent of growth during each of these growth phases.

The size of budding yeast cells is further influenced by growth rate. For example, yeast cells growing slowly in poor nutrients are nearly half the size of cells in rich nutrients (Johnston *et al*., 1977; 1979). This observation illustrates a peculiar and poorly understood aspect of growth control: cell size is proportional to growth rate (Ferrezuelo *et al*., 2012; Leitao and Kellogg, 2017). The relationship holds when comparing cells growing under different nutrient conditions that support different growth rates, and when comparing cells that show different growth rates despite growing under identical nutrient conditions. Conversely, growth rate is proportional to cell size in yeast (Elliott and McLaughlin, 1978; Schmoller et al., 2015; Leitao and Kellogg, 2017). For example, the growth rate of the daughter bud is proportional to the size of the mother cell. A proportional relationship between cell size and growth rate appears to hold across all orders of life (Schaechter *et al*., 1958; Robertson, 1963; Hirsch and Han, 1969; Fantes and Nurse, 1977; Johnston *et al*., 1977).

Clues to a mechanistic basis for the relationship between cell size and growth rate in budding yeast have come from analysis of a form of protein phosphatase 2A that is bound to the Rts1 regulatory subunit (PP2A^Rts1^). PP2A^Rts1^ is required for normal control of cell growth and size (Artiles *et al*., 2009; Zapata *et al*., 2014). Thus, cells that lack PP2A^Rts1^ are abnormally large and fail to reduce their size in poor nutrients. PP2A^Rts1^ relays signals that influence a TORC2 signaling network that is required for normal control of cell size and growth rate (Lucena *et al*., 2017). The TORC2 network controls synthesis of ceramide lipids, which play roles in signaling. Genetic and pharmacological data suggest that modulation of ceramide synthesis by PP2A^Rts1^ and TORC2 is required for nutrient modulation of cell size and growth rate in G1 phase. Cells that can not synthesize ceramides fail to increase their growth rate during G1 phase when shifted from poor to rich carbon, and they complete G1 phase at identical small sizes in rich or poor carbon. It is unknown whether signals from the PP2A^Rts1^ network also control cell growth and size during the mitotic growth interval.

Here, we have investigated control of cell growth and size during growth in mitosis. Cell size at completion of mitosis is proportional to growth rate during mitosis (Leitao and Kellogg, 2017). Moreover, when growth rate is slowed by poor nutrients, the duration of mitosis is increased, which likely reflects mechanisms that ensure that completion of mitotic events occurs only when a sufficient amount of growth has occurred (Leitao and Kellogg, 2017). Together, these observations suggest that growth during mitosis is regulated. A proteome-wide mass spectrometry search for proteins controlled by PP2A^Rts1^ identified numerous proteins implicated in control of mitotic progression and cell size, which suggested that PP2A^Rts1^ could play an important role in control of cell growth and size in mitosis (Zapata *et al*., 2014). Thus, as a first step we tested whether PP2A^Rts1^ influences cell growth and size in mitosis.

## Results and Discussion

### PP2A^Rts1^ is required for the proportional relationship between cell size and growth rate

We first tested how inactivation of PP2A^Rts1^ influences mitotic duration and daughter cell size. Previous studies have shown that mother cell size strongly influences daughter cell size and growth rate (Johnston et al., 1977; Schmoller et al., 2015; Leitao and Kellogg, 2017). Therefore, interpretation of results from *rts1*Δ cells would be complicated by the possibility that effects on bud growth and size are a secondary consequence of defects in mother cell size that arose in previous generations. To avoid this problem, we fused *RTS1* to an auxin-inducible degron (*rts1-AID*), which allowed us to observe the immediate effects of inactivating PP2A^Rts1^ (Nishimura *et al*., 2009). In the absence of auxin, *rts1-AID* did not cause obvious defects in cell size or cell cycle progression (Figures S1A,B). Addition of auxin caused rapid destruction of rts1-AID protein within 15–30 minutes (Figure S1C). Approximately 10% of the rts1-AID protein remained in the presence of auxin. In addition, *rts1-AID* cells growing in the presence of auxin at elevated temperatures formed colonies more rapidly than *rts1*Δ cells (Figure S1D). Together, these observations suggest that *rts1-AID* causes a partial loss of function when auxin is present. Nevertheless, we utilized *rts1-AID* because it allowed analysis of bud growth and mitotic duration without the complication of aberrant mother cell size.

We used a microscopy-based assay (Ferrezuelo et al., 2012; Leitao and Kellogg, 2017) to test how loss of *rts1-AID* affected growth during mitosis. Auxin was added to synchronized cells shortly after bud emergence. Destruction of rts1-AID caused a 3-fold increase in the average duration of metaphase and anaphase in both rich and poor carbon (Figure 1A; see Figure S2A for dot plots and p-values). Destruction of rts1-AID also caused a large increase in the variance of metaphase duration compared to wild type cells (Figure S2A).

**Figure 1.**
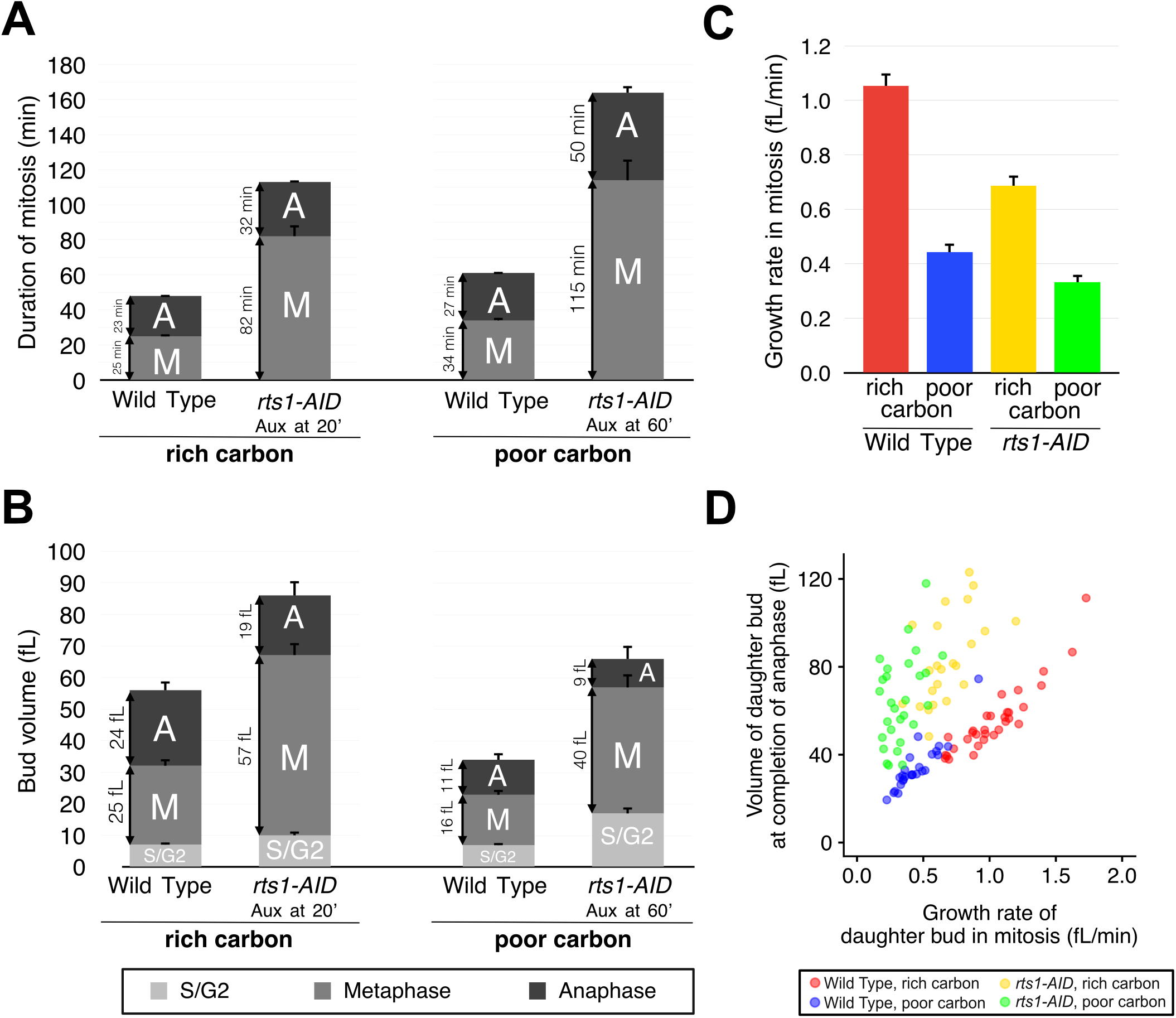
PP2A^Rts1^ is required for normal control of mitotic duration and cell size. (**A**) Plots showing the average durations of metaphase and anaphase for wild type and *rts1-AID* cells growing in rich or poor carbon. (**B**) The average growth in volume for all phases of the cell cycle except G1 phase is plotted for wild type and *rts1-AID* cells growing in rich or poor carbon. Due to the extended length of the cell cycle in *rts1-AID* cells, only a few cells could be followed through G1. For this reason, the limited data regarding G1 phase were omitted. (**C**) The growth rate during metaphase and anaphase was calculated as the average of individual cell growth rates. (**D**) A plot of the volume of the daughter bud at completion of anaphase versus growth rate during metaphase plus anaphase. Error bars represent standard error of the mean.

We next analyzed daughter bud size at each mitotic transition. *rts1-AID* caused a large increase in daughter bud size at completion of each stage of mitosis in both rich and poor carbon (Figure 1B; see Figures S2C,D for dot plots and p-values). The variance in size at the end of metaphase was much larger in *rts1-AID* cells compared to wild type cells. Importantly, there was not a statistically significant difference in the size of *rts1-AID* daughter cells in rich or poor carbon at the end of metaphase (Figure S2C). By the end of anaphase, *rts1-AID* cells in rich carbon were slightly larger than their counterparts in poor carbon (Figure S2D).

**Figure 2.**
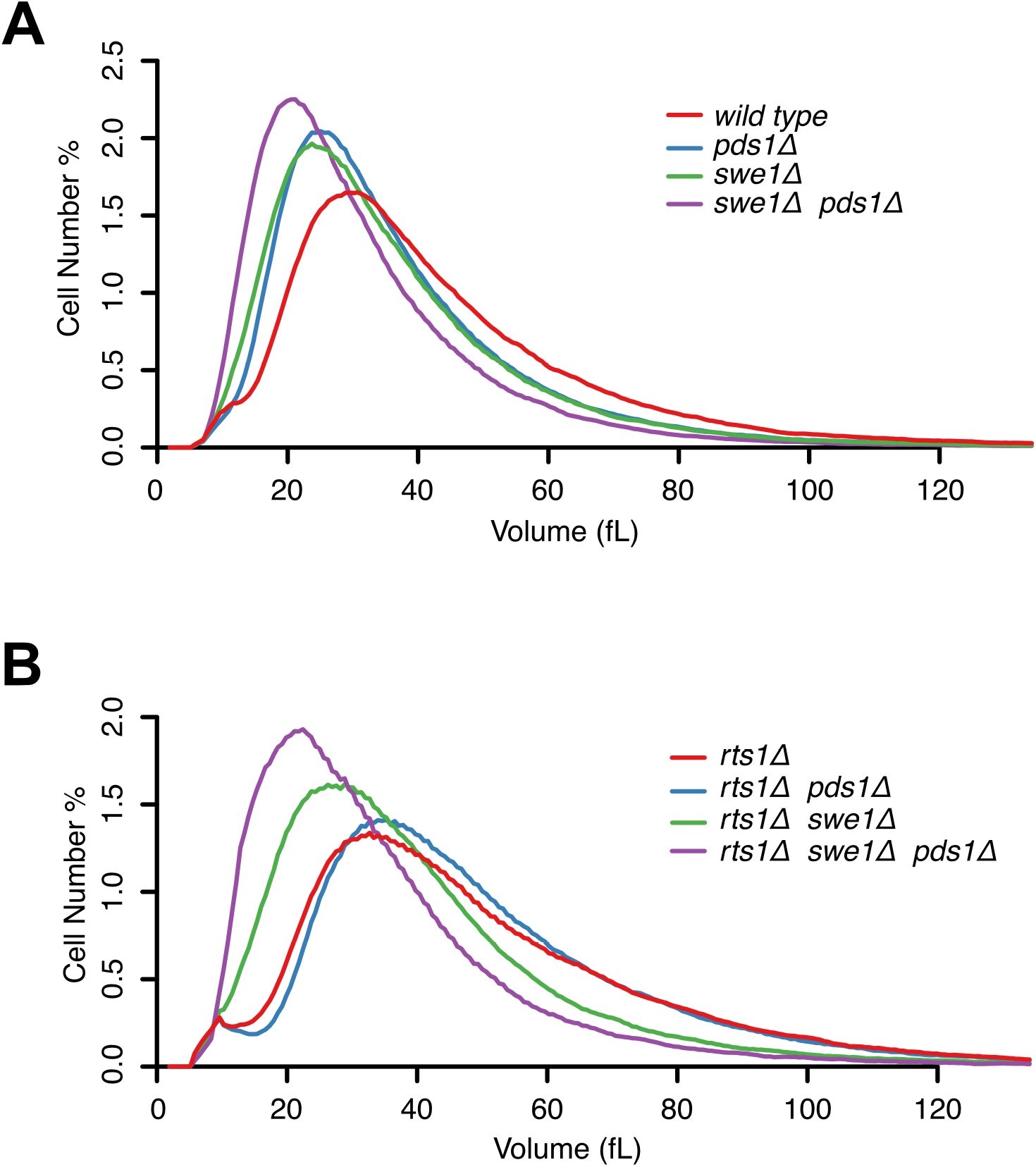
Swe1 and Pds1 influence cell size. (**A**,**B**) Size distributions of log phase populations of cells of the indicated genotypes were measured with a Coulter Counter. Cells were grown in YPD medium.

Further analysis of the *rts1-AID* allele led to surprising insight into the role of PP2A^Rts1^ in cell size control. Destruction of rts1-AID reduced the average rate of growth in mitosis by approximately 30% in both rich and poor carbon (Figure 1C). In normal cells, reduced growth rate causes reduced cell size; however, destruction of rts1-AID caused an increase in size. In addition, the correlation between growth rate and cell size was broken in *rts1-AID* cells (Figure 1D). Thus, buds with nearly identical growth rates completed mitosis at very different sizes. Conversely, cells that completed mitosis at identical sizes grew at very different rates during mitosis. A previous study found that inactivation of Ydj1, a member of the JJJ chaperone family, causes a similar discordance between cell size and growth rate at the end of G1 phase (Ferrezuelo *et al*., 2012). Proteome-wide mass spectrometry analysis of *rts1*Δ cells suggests that Ydj1 is controlled by PP2A^Rts1^ (Zapata *et al*., 2014).

To summarize, a partial loss of PP2A^Rts1^ function caused major defects in cell size control in mitosis, as well as a nearly complete loss of the proportional relationship between cell size and growth rate. These observations are consistent with our previous discovery that *rts1*Δ causes a nearly complete failure in modulation of cell size in response to changes in carbon source (Artiles *et al*., 2009).

As a next step, we sought to identify proteins that respond to PP2A^Rts1^-dependent signals and control the duration of growth in mitosis. Over the long term, identification of these proteins and the signals that control their activity should yield insight into mechanisms that control cell growth and size.

### Swe1 and Pds1 are required for normal control of cell size at completion of mitosis

We used a candidate approach to identify proteins that control the duration of growth in mitosis. To identify candidates, we used two criteria. First, we considered proteins that were previously found to control cell size and/or the duration of mitosis. Second, we considered potential targets of PP2A^Rts1^-dependent regulation that were previously identified by proteome-wide mass spectrometry analysis of *rts1*Δ cells, or by phenotypic analysis of *rts1*Δ cells (Harvey *et al*., 2011; Zapata *et al*., 2014). Two candidates fulfilled both criteria: Swe1 and Pds1/securin.

Swe1 is the budding yeast homolog of the Wee1 kinase, which delays mitosis by phosphorylating and inhibiting mitotic Cdk1. Wee1 was originally found to control mitotic entry. However, more recent studies found that Wee1 family members also control the duration of mitosis in both yeast and human cells (Jin *et al*., 2008; Raspelli *et al*., 2011; Lianga *et al*., 2013; Toledo *et al*., 2015). Moreover, mass spectrometry identified the inhibitory site on Cdk1 targeted by Swe1 as the most strongly hyperphosphorylated site in *rts1*Δ cells (Zapata *et al*., 2014). The mass spectrometry also showed that Swe1 is hyperphosphorylated in *rts1*Δ cells on multiple sites that play a role in its activation, and analysis of Swe1 phosphorylation in *rts1*Δ cells by western blotting has shown that Swe1 persists in a partially hyperphosphorylated form that is thought to be the active form (Harvey *et al*., 2011). Finally, there is evidence that PP2A^Rts1^ can also control removal of Cdk1 inhibitory phosphorylation independently of its role in controlling Swe1 phosphorylation (Kennedy *et al*., 2016). Together, these observations demonstrate that inhibitory phosphorylation of Cdk1 could play a role in prolonging mitosis when growth is slowed in poor carbon.

The second candidate, Pds1/securin, inhibits chromosome segregation by binding and inhibiting separase, a protease that cleaves cohesins that hold chromosomes together (Cohen-Fix and Koshland, 1997; Ciosk *et al*., 1998; Uhlmann *et al*., 1999). Exit from mitosis is triggered by activation of the anaphase-promoting complex (APC), which targets Pds1 for destruction. Pds1 and separase also control mitotic cyclin destruction, which indicates that they can control the duration of mitosis (Cohen-Fix and Koshland, 1999; Tinker-Kulberg and Morgan, 1999). In addition, DNA damage induces a mitotic arrest by triggering signals that lead to phosphorylation of Pds1, thereby protecting it from the APC (Yamamoto *et al*., 1996; Wang *et al*., 2001). Thus, there is a precedent for signals upstream of Pds1 controlling the duration of mitosis. Finally, proteome-wide mass spectrometry data suggest that Pds1 is hyperphosphorylated in *rts1*Δ cells (Zapata *et al*., 2014).

If Pds1 and Swe1 play roles in prolonging mitosis to allow growth to occur, then their loss should cause reduced cell size. Previous studies have shown that *swe1*Δ reduces cell size, but there has been no evidence that Pds1 is required for cell size control (Jorgensen *et al*., 2002; Harvey and Kellogg, 2003; Harvey *et al*., 2005; 2011). Coulter counter analysis of cell size revealed that *pds1*Δ caused a reduction in cell size similar to the reduction in cell size caused by *swe1*Δ (Figure 2A). When combined, *swe1*Δ and *pds1*Δ caused an additive decrease in cell size.

We next hypothesized that the prolonged duration of mitosis and increased cell size caused by loss of PP2A^Rts1^ is due to hyperactivity of Swe1 and Pds1. To begin to test this, we first analyzed the effects of *swe1*Δ and *pds1*Δ on the size of *rts1*Δ cells. The abnormally large size of *rts1*Δ cells was eliminated by *swe1Δ pds1*Δ, which suggested that the increased duration of growth in mitosis in *rts1-AID* cells could be due entirely to hyperactive Pds1 and Swe1 (Figure 2B). *pds1*Δ alone did not reduce the size of *rts1*Δ cells. We suspect that this is the result of hyperactive Swe1 in *rts1*Δ cells, which would lead to decreased Cdk1 activity. Since Cdk1 is required for activation of the anaphase promoting complex, this could lead to a mitotic delay.

### Inhibitory phosphorylation of Cdk1 is partially responsible for the increased duration of mitosis in poor carbon

We next investigated the role of Cdk1 inhibitory phosphorylation more closely. To test whether Swe1 plays a role in prolonging mitosis in poor carbon, we first tested whether Cdk1 inhibitory phosphorylation is prolonged in poor carbon. Cells growing in rich or poor carbon were released from a G1 arrest and Cdk1 inhibitory phosphorylation was analyzed with a phosphospecific antibody. Samples were also analyzed by immunofluorescence to determine the fraction of cells with metaphase spindles at each time point. Cdk1 inhibitory phosphorylation was prolonged in poor carbon and was correlated with the presence of metaphase spindles (Figure 3A).

**Figure 3.**
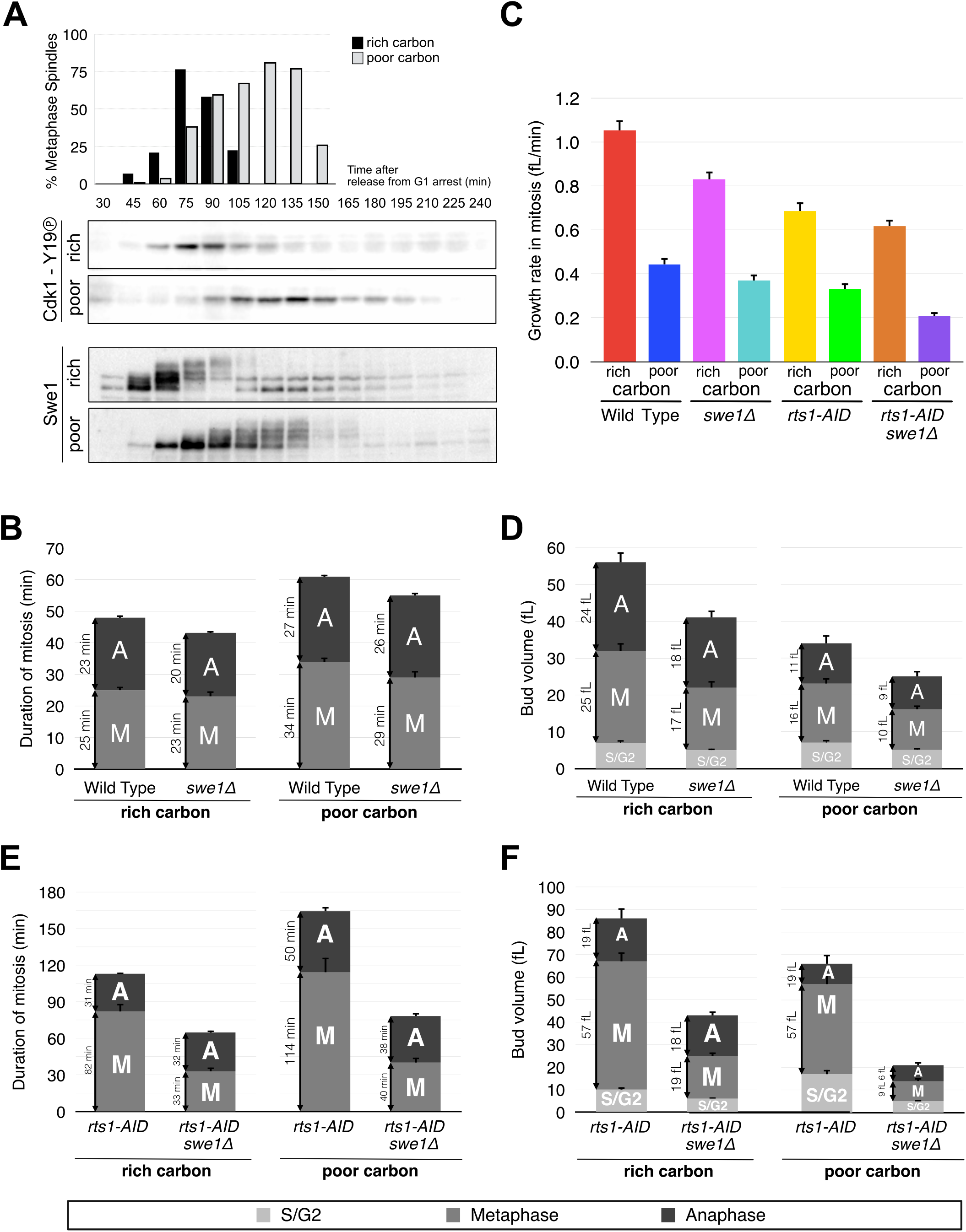
The increased duration of mitosis in poor carbon is partially due to Cdk1 inhibitory phosphorylation. (**A**) Wild type cells growing in rich or poor carbon were released from a G1 arrest and samples were taken at 15 minute intervals. The percentage of cells with a metaphase spindle was determined by immunofluorescence microscopy. Cdk1 inhibitory phosphorylation on tyrosine 19 was detected by western blot with a phosphospecific antibody. Swe1 was detected by western blot with an anti-Swe1 antibody. Cdk1 inhibitory phosphorylation and Swe1 were analyzed in the same samples. Mitotic spindle data are from an independent biological replicate that showed similar timing of events. (**B**) Average durations of metaphase and anaphase for wild type and *swe1*Δ cells growing in rich or poor carbon. (**C**) Average growth rates during metaphase and anaphase for wild type and *swe1*Δ cells in rich or poor carbon. (**D**) Average growth in volume during metaphase and anaphase for wild type or *swe1*Δ cells growing in rich or poor carbon. (**E**) Average durations of metaphase and anaphase for *rts1-AID* and *rts1-AID swe1*Δ cells growing in rich or poor carbon. (**F**) Average growth in volume during metaphase and anaphase for *rts1-AID* and *rts1-AID swe1*Δ cells growing in rich or poor carbon. Error bars represent standard error of the mean.

We also analyzed Swe1 phosphorylation during the cell cycle in rich and poor carbon. Swe1 passes through multiple phosphorylation states during mitosis that can be detected via electrophoretic mobility shifts; attainment of a fully hyperphosphorylated state is correlated with inactivation of Swe1 (Sreenivasan and Kellogg, 1999; Harvey *et al*., 2005; 2011). In rich media, Swe1 reached full hyperphosphorylation and was degraded shortly thereafter (Figure 3A). In poor media, Swe1 was present throughout much of the prolonged mitosis. Moreover, Swe1 took longer to reach the fully hyperphosphorylated state, and it persisted in the partially hyperphosphorylated state that is thought to represent active Swe1 (Harvey *et al*., 2011). These observations suggest that signals that control Swe1 could prolong Cdk1 inhibitory phosphorylation in poor nutrients.

The role of Swe1 was further characterized by analyzing daughter cell growth and mitotic events in single cells. In rich carbon, *swe1*Δ caused a slight reduction in the average duration of metaphase, as previously described (Figure 3B; see Figure S2A for dot plots and p-values) (Lianga *et al*., 2013). In poor carbon, *swe1*Δ caused a greater reduction in metaphase duration, from 34 minutes to 29 minutes. Although *swe1*Δ reduced the duration of metaphase in poor carbon, it did not reduce it to the mitotic duration observed for wild type cells in rich carbon, which indicated that the mitotic delay caused by poor carbon is not due solely to inhibitory phosphorylation of Cdk1. Loss of Swe1 had little effect on the duration of anaphase, consistent with the observation that Cdk1 inhibitory phosphorylation is observed primarily during metaphase (Figure 3A).

In both rich and poor carbon, *swe1*Δ caused a reduction in growth rate (Figure 3C). This, combined with the reduced duration of metaphase, caused *swe1*Δ daughter buds to undergo mitotic transitions at a substantially reduced size in both rich and poor carbon (Figure 3D; see Figures S2C,D for dot plots and p-values). Together, the data demonstrate that Swe1 plays a role in the increased duration of metaphase in poor carbon and is required for normal control of daughter cell size at cytokinesis. The reduced growth rate of daughter buds in *swe1*Δ cells could be due to the reduced size of their mother cells.

We next defined the contribution of Cdk1 inhibitory phosphorylation to the mitotic delay observed in *rts1-AID* cells. Western blot assays confirmed that destruction of rts1-AID caused a prolonged mitotic delay in both rich and poor carbon (Figure S3A). The delay caused by *rts1-AID* in rich carbon was reduced, but not eliminated, by *swe1*Δ. We further discovered that *rts1*Δ caused a mitotic delay after release from a metaphase arrest. The delay was reduced by *swe1*Δ, but not fully eliminated (Figure S3B).

In single cell assays, the metaphase delays caused by *rts1-AID* in rich and poor carbon were reduced by *swe1*Δ, but not fully eliminated (Figure 3E; see Figure S2A for dot plots and p-values). The increased duration of anaphase caused by *rts1-AID* in rich and poor carbon was largely unaffected by *swe1*Δ. Although *swe1*Δ did not fully rescue the mitotic delays caused by *rts1-AID*, it caused *rts1-AID* cells to exit mitosis at sizes similar to those of *swe1*Δ cells (Figure 3F; see Figure S2 for dot plots and p-values). This was a combined result of the reduction in mitotic duration and decreased growth rate in *rts1-AID swe1*Δ cells relative to *rts1-AID* or *swe1*Δ cells.

Together, these observations demonstrate that nutrients and PP2A^Rts1^ control mitotic duration and daughter cell size via a Swe1-dependent pathway, as well as by a Swe1-independent pathway. A previous study found that purified PP2A^Rts1^ can not dephosphorylate

Swe1 in vitro, so it is likely that it controls Swe1 indirectly (Harvey *et al*., 2011).

### Phosphorylation of Pds1 is controlled by nutrients and PP2A^Rts1^

We next investigated the function and regulation of Pds1. Proteome-wide mass spectrometry analysis identified four serines in Pds1 that are hyperphosphorylated in *rts1*Δ cells (S185, S186, S212, S213) (Zapata *et al*., 2014). Two of the sites (S212,S213) were previously implicated in delaying mitosis in response to DNA damage (Wang *et al*., 2001). We hypothesized that hyperphosphorylation of Pds1 at these sites delays mitotic progression in poor carbon and in *rts1*Δ cells. To test this, we first used Phos-Tag gels to determine whether Pds1 undergoes hyperphosphorylation when cells are shifted to poor carbon. Hyperphosphorylated forms of Pds1 could be detected within 5 minutes of a shift to poor carbon (Figure 4A, lanes 1–3). Moreover, *rts1*Δ caused Pds1 to undergo more rapid and extensive hyperphosphorylation in response to poor carbon (Figure 4A, lanes 5–7). We also observed that *rts1*Δ caused Pds1 to undergo a slight shift to hyperphosphorylated forms in cells growing in rich carbon (Figure 4A, compare lanes 1 and 5). A mutant version of Pds1 that lacks the 4 sites controlled by PP2A^Rts1^ (*pds1–4A*) largely failed to undergo hyperphosphorylation when shifted to poor carbon (Figure 4A, lanes 9–11). Thus, Pds1 undergoes rapid phosphorylation in response to carbon source, and PP2A^Rts1^ is required for control of Pds1 phosphorylation.

**Figure 4.**
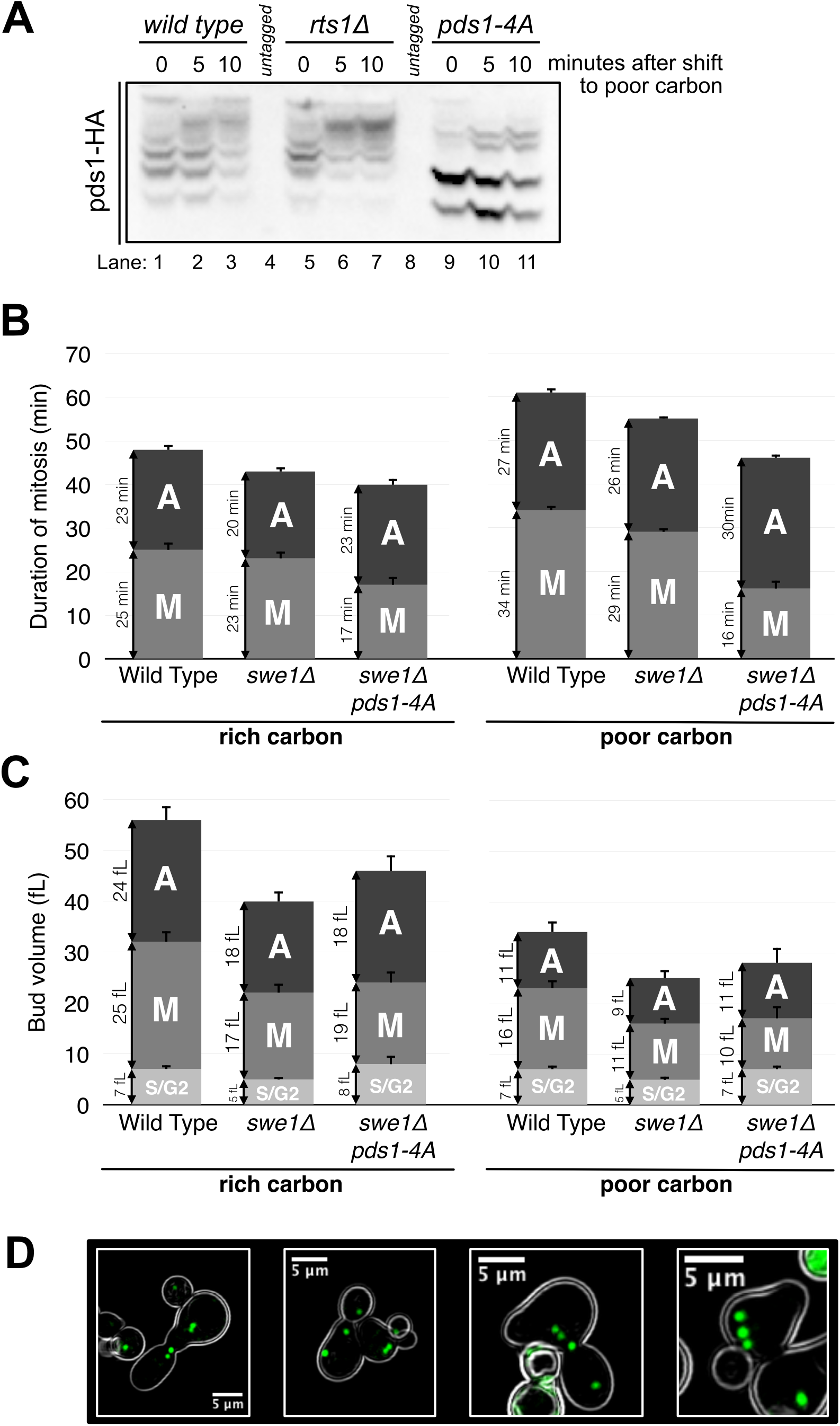
Pds1 and Swe1 are required for nutrient modulation of the duration of metaphase. (**A**) Wild type, *rts1*Δ and *pds1–4A* cells grown to log phase were switched from YPD to YPG/E and Pds1 phosphorylation was assayed by PhosTag western blot. (**B**) Average durations of metaphase and anaphase for wild type, *swe1*Δ and *swe1*Δ *pds1–4A* cells in rich or poor carbon. (**C**) Average growth in volume for all phases of the cell cycle except G1 phase is plotted for wild type, *swe1*Δ and *swe1*Δ *pds1–4A* cells in rich or poor carbon. (**D**) Examples of *swe1*Δ *pds1–4A* cells with multiple spindle poles.

### Pds1 and Swe1 are required for nutrient modulation of the duration of metaphase

To test whether Pds1 contributes to the mitotic delay observed in *swe1*Δ cells in poor carbon, we analyzed the effects of *pds1–4A* on the duration of mitosis in *swe1*Δ cells. Metaphase in *swe1Δ pds1–4A* cells was shorter than metaphase in *wild type* cells or *swe1*Δ cells in both rich and poor carbon (Figure 4B; see Figure S2A for dot plots and p-values). In addition, there was no difference in the duration of metaphase between rich and poor carbon in the *swe1Δ pds1–4A* cells. Importantly, these data suggest that nutrient modulation of the duration of metaphase could be due solely to regulation of Swe1 and Pds1.

*pds1–4A swe1*Δ cells growing in poor carbon completed metaphase at a reduced volume compared to *pds1–4A swe1*Δ cells growing in rich carbon (Figure 4C; see Figure S2C for dot plots and p-values). This was entirely a consequence of reduced growth rate in poor carbon, since the duration of metaphase in *pds1–4A swe1*Δ cells was identical in rich and poor carbon. Thus, *pds1–4A swe1*Δ causes cell size at completion of metaphase to become a simple function of growth rate, as originally imagined in early models of cell size control (Hartwell and Unger, 1977).

In both rich and poor carbon, the duration of anaphase and the extent of growth during anaphase were increased in *pds1–4A swe1*Δ cells relative to *swe1*Δ cells. In addition, the duration of anaphase in the *pds1–4A swe1*Δ cells was modulated by nutrients. These observations suggest that the mechanisms that control the duration of anaphase in response to carbon source are largely independent of Swe1 and Pds1. The increased duration of anaphase in *pds1–4A swe1*Δ cells may reflect compensatory growth that occurs because the cells complete metaphase at an abnormally small size. Proteome-wide mass spectrometry identified numerous components of the mitotic exit network as potential targets of PP2A^Rts1^-dependent regulation (Zapata *et al*., 2014). Thus, nutrient-dependent control of the mitotic exit network could account for the increased duration of growth in anaphase in poor nutrients.

The average size of *pds1–4A swe1*Δ buds at completion of anaphase in poor carbon was slightly larger than *swe1*Δ cells, which was due to a few very large outlier cells (see dot plots in Figure S2D). We noticed that some *pds1–4A swe1*Δ cells in poor carbon had more than two spindle pole bodies (Figure 4D). This suggested that they have multiple nuclei, which could be a consequence of a failure to undergo sufficient growth before nuclear division. Cells in which we could detect extra spindle poles, which constituted approximately 10% of the cells growing in poor carbon, were excluded from the analysis in Figures 4B and 4C. However, the large outlier cells in the *pds1–4A swe1*Δ data could be polyploid cells that did not have well separated spindle poles that would clearly identify them as having multiple nuclei. Since cell size scales with ploidy, the unusually large *pds1–4A swe1*Δ cells could correspond to polyploid cells.

### Pds1 and Swe1 are not required for the proportional relationship between cell size and growth rate

A plot of cell size at completion of mitosis versus growth rate revealed that *pds1–4A swe1*Δ cells have a reduced size, yet show a proportional relationship between cell size and growth rate (Figure 5A). Moreover, analysis of *pds1Δ swe1*Δ cell size distributions showed that they are abnormally small, yet still show evidence of nutrient modulation of cell size (Figure 5B). In contrast, *rts1Δ pds1Δ swe1*Δ cells show little evidence of nutrient modulation of cell size (Figure 5C).

**Figure 5.**
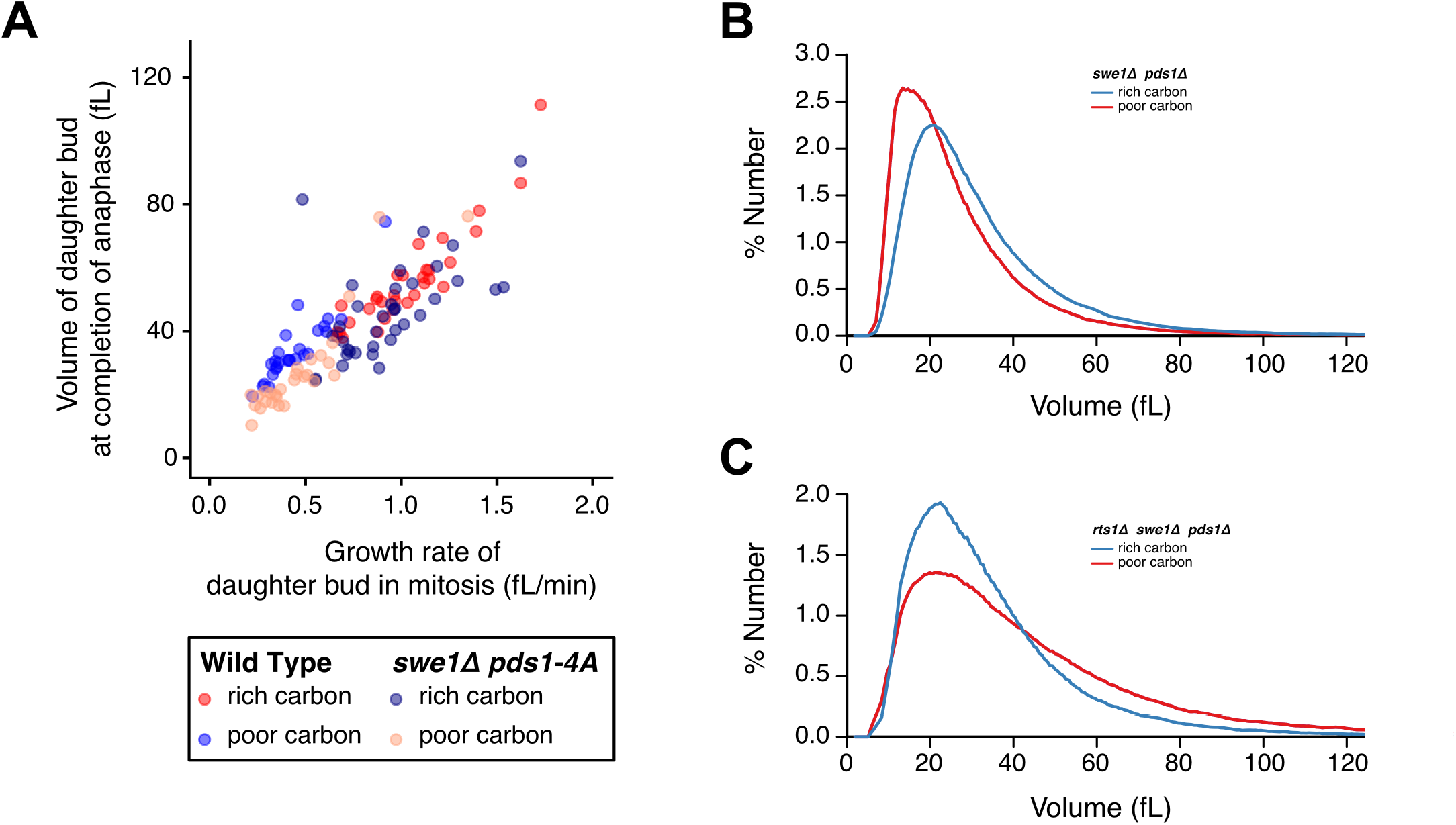
Pds1 and Swe1 are not required for the proportional relationship between cell size and growth rate. (**A**) A plot of the volume of the daughter bud at completion of anaphase versus growth rate during metaphase plus anaphase. (**B,C**) Size distributions of log phase populations of cells of the indicated genotypes were measured with a Coulter Counter. Cells were grown in YPD medium.

Together, the data show that Pds1 and Swe1, by virtue of their ability to increase the duration of growth in mitosis, strongly influence how nutrients modulate cell size. Thus, cells that lack Pds1 and Swe1 are abnormally small and fail to undergo an appropriate increase in size when shifted from poor to rich carbon. The data further suggest that Pds1 and Swe1 are controlled by PP2A^Rts1^ and nutrient-dependent signals, consistent with a role in setting cell size in response to growth rate. Yet Pds1 and Swe1 are not strictly required for the proportional relationship between cell size and growth rate during growth in mitosis, whereas PP2A^Rts1^ is required.

In previous work, we found that PP2A^Rts1^ influences a TORC2 signaling network that is required for normal control of cell growth and size in G1 phase (Lucena *et al*., 2017). The TORC2 network includes a feedback loop in which TORC2 promotes production of ceramides, while ceramides generate signals that inhibit TORC2 signaling. An inhibitor of ceramide production causes a dose-dependent decrease in growth rate during G1 phase, as well as a decrease in cell size. Thus, manipulation of signals within the TORC2 network in cells that have functional PP2A^Rts1^ causes strong effects on growth rate and cell size, yet the proportional relationship between cell size and growth rate appears to be maintained.

Here, we have extended these studies to show that PP2A^Rts1^ strongly influences cell growth and size in mitosis. Importantly, we also show that PP2A^Rts1^ is required for a proportional relationship between cell size and growth rate during mitosis. To explain the data, we suggest that PP2A^Rts1^ relays signals within the TORC2 network that ensure a proportional relationship between cell size and growth rate. Cell growth is driven by complex signaling networks that control the rates of the diverse pathways that comprise cell growth. Since the rate of growth in mitosis is proportional to cell size, one might expect that the signals that drive growth are also proportional to cell size, and that specific mechanisms ensure that the signals that drive growth rate scale with size. In this case, a failure in a PP2A^Rts1^-dependent mechanism that makes growth rate proportional to cell size should cause cells to grow at rates that are uncorrelated with size, leading to a loss of the correlation between cell size and growth rate. The correlation would breakdown in both directions – growth rate would no longer be proportional to cell size, and cell size would no longer be proportional to growth rate because cells of the same size would have different growth rates. The fact that PP2A^Rts1^ influences signaling within the TORC2 feedback loop suggests that it is well-positioned to enforce a mechanistic link between cell size and growth rate (Lucena *et al*., 2017). Moreover, there is evidence that the level of signaling in the feedback loop is influenced by growth, which might be expected for a signaling network that ensures that cell size and growth rate are proportional (Clarke *et al*., 2017). For example, one way to enforce a proportional relationship between cell size and growth rate would be to have the events of growth generate feedback signals that modulate the signals that drive growth.

The data further suggest that the PP2A^Rts1^-dependent signals that make cell size proportional to growth rate also control the activity of Pds1 and Swe1 to set the threshold amount of growth required for cell cycle progression. In this view, Pds1 and Swe1 would mediate PP2A^Rts1^-dependent signals that delay cell cycle progression until growth reaches a threshold appropriate for the growth rate. Thus, Pds1 and Swe1 would be required for cells to reach a size that is appropriate for the growth rate, but they would not be required for the proportional relationship between cell size and growth rate.

A model in which the same global signals that set growth rate also set the critical amount of growth required for cell cycle progression would provide a simple mechanistic explanation for the proportional relationship between cell size and growth rate. Since cell size control likely evolved as an outcome of growth control, it would make sense that control of cell growth and size are mechanistically linked. Further analysis of the signals that connect Pds1 and Swe1 to PP2A^Rts1^ and the TORC2 network should lead to important clues to how cell growth and size are controlled.

### Materials and Methods

#### Yeast strains and media

The genotypes of the strains used in this study are listed in Table 1. All strains are in the W303 background (*leu2–3,112 ura3–1 can1–100 ade2–1 his3–11,15 trp1–1 GAL+, ssd1-d2*). Genetic alterations were carried out using one-step PCR-based integration at the endogenous locus (Longtine *et al*., 1998; Janke *et al*., 2004) or by genetic crossing.

**Table 1.**

| Strain | Mating type | Genotype |
| --- | --- | --- |
| DK186 | a | *bar1* |
| DK2423 | a | *bar1 TIR1::LEU2 TIR1::HIS3 rts1-AID::KanMX6* |
| DK2523 | a | *bar1 SPC42-GFP::HIS3 MYO1-GFP::TRP* |
| DK2879 | a | *bar1 TIR1::LEU2 TIR1::HIS3 SPC42-GFP::hphNT1 MYO1-GFP::KITRP rts1-AID::KanMX6* |
| DK2930 | a | *bar1 SPC42-GFP::HIS3, MYO1-GFP::TRP swe1Δ::URA3* |
| DK3072 | a | *bar1 TIR1::LEU2, TIR1::HIS3, SPC42-GFP::hphNT1 MYO1-GFP::KITRP rts1-AID::KanMX6 swe1Δ::URA3* |
| DK1993 | a | *bar1 GAL1-CDC20::NatNT2* |
| DK2176 | a | *bar1 GAL1-CDC20::NatNT2 rts1Δ::KanMX4* |
| DK2243 | a | *bar1 GAL1-CDC20::NatNT2 rts1Δ::KanMX4 swe1::URA3* |
| DK1140 | α | *bar1 rts1Δ::HIS3* |
| DK647 | a | *bar1 rts1Δ::KANMX4* |
| DK2721 | a | *bar1 pds1Δ::LEU2* |
| SH24 | a | *bar1 swe1Δ::URA3* |
| **DK2722** | a | *bar1 pds1Δ::LEU2 swe1Δ::URA3* |
| **DK3363** | a | *bar1 rts1Δ::HIS pds1Δ::LEU2* |
| **DK3365** | a | *bar1 rts1::HIS swe1::URA3* |
| **DK3361** | a | *bar1 rts1Δ::HIS swe1Δ::URA3 pds1Δ::LEU2* |
| **DK1778** | a | *bar1 PDS1-3XHA::TRP* |
| **DK2678** | a | *bar1 PDS1-3XHA::TRP rts1Δ::KanMX4* |
| **DK3318** | a | *bar1 pds1-4A-3XHA::TRP* |
| **DK815** | α | *bar1 swe1Δ::URA3 rts1Δ::KanMX4* |
| **DK3335** | a | *bar1 pds1-4A-3XHA::TRP swe1Δ::URA3 SPC42-GFP::HIS3* |

*pds1–4A* mutants were created by replacing a 170bp fragment of *PDS1* that contains S185, S186, S212 and S213 with the *URA3* marker. The replacement was carried out in a *PDS1–3xHA::TRP* strain so that the final mutant version of *PDS1* would be tagged with 3XHA. The *URA3* marker gene was replaced by a mutant fragment of *PDS1* in which all 4 sites were mutated to alanine, which was synthesized by overlap PCR. The mutant fragment was co-transformed with a *LEU2* marked *CEN* vector (YCplac111) and LEU+ transformants were selected to enrich for transformation-competent cells. The *LEU+* transformants were then replica plated to FOA to select for cells that lost the *URA3* marker. Pre-selection for the *LEU2 CEN* vector dramatically reduced the background of spontaneous *ura3* mutants. The resulting strain was backcrossed once to wild type and then to DK815 to obtain DK3320, which was transformed with SPC42-GFP::HIS to obtain DK3335.

For cell cycle time courses and analysis of cell size by Coulter counter, cells were grown in YP media (1% yeast extract, 2% peptone, 8ml/L adenine) supplemented with 2% dextrose (YPD), or with 2% glycerol and 2% ethanol (YPGE). For microscopy, cells were grown in complete synthetic media (CSM) supplemented with 2% dextrose (CSM-Dex) or 2% glycerol and 2% ethanol (CSM-G/E).

#### Microscopy and image analysis

Microscopy, image analysis, and statistical analysis of microscopy data were carried out as previously described (Leitao and Kellogg, 2017)

#### Cell cycle time courses, western blotting and Coulter counter analysis

Cell cycle time courses utilizing cells arrested in G1 phase were carried out as previously described. To arrest cells in mitosis, *GAL1-CDC20* cells were grown overnight in YP media containing 2% raffinose and 2% galactose and arrested by washing into YP containing 2% raffinose. Cells were monitored until most cells had large buds. Cells were released from metaphase by re-addition of 2% galactose. SDS-polyacrylamide gel electrophoresis and western blotting were carried out as previously described. Blots were probed overnight at 4°C with affinity-purified rabbit polyclonal antibodies raised against Swe1 or with a phosphospecific antibody that recognizes Cdk1 phosphorylated at tyrosine 19 (Cell Signaling Technology, cat# 10A11). Blots were exposed to film or imaged using a ChemiDoc™ MP System (Bio-Rad). For quantification of rts1-AID degradation, band intensity was quantified using ImageLab^TM^. Destruction of Rts1-AID was initiated by addition of 1 mM auxin from a 50 mM stock made in 100% ethanol.

For PhosTag western blots, cells were lysed by bead-beating into sample buffer without phosphatase inhibitors. After cell lysis, samples were centrifuged for 1 min at 13,000 rpm at 4°C and quickly placed in a boiling water bath for 7 min. Samples were loaded into 10% SDS-polyacrylamide gels supplemented with 100 µM PhosTag and 200 μM MnCl_2_. To prepare PhosTag gels, the gel mixture was degassed for 5 min prior to addition of TEMED and polymerization was allowed to occur for 1–2 hours at room temperature followed by overnight at 4°C. Gels were run at 2.5 mA for 16 hrs until a 29-kD marker was at the bottom of the gel. The gel was incubated for 10 min in transfer buffer supplemented with 2 mM EDTA, followed by a second incubation without EDTA. Gels were transferred onto nitrocellulose via the Trans-Blot Turbo Transfer System (Bio-Rad). Blots were probed at room temperature with the 12CA5 anti-HA monoclonal antibody followed by HRP-conjugated anti-mouse secondary antibody. Secondary antibodies were detected via chemiluminescence using Quantum substrate (Advansta).

Analysis of cell size by Coulter counter was carried out as previously described (Leitao and Kellogg, 2017).

## Acknowledgments

We thank Ben Abrams, Facilities Manager for the UCSC Life Sciences Microscopy Center for support and mentoring with all microscopy related techniques, and the Aldea lab for sharing BudJ: an ImageJ plugin to analyze images of budding yeast cells (http://www.ibmb.csic.es/home/maldea). R.L. was supported by a fellowship from the “Fundação para a Ciência e a Tecnologia” (FCT), with funds from “Programa Operacional Potencial Humano/ Fundo Social Europeu” (POPH/FSE), under the fellowship SFRH/BD/75004/2010. This work was supported by National Institutes of Health Grant R01-GM053959.

## Supplemental Figure Legends

**Figure S1:**
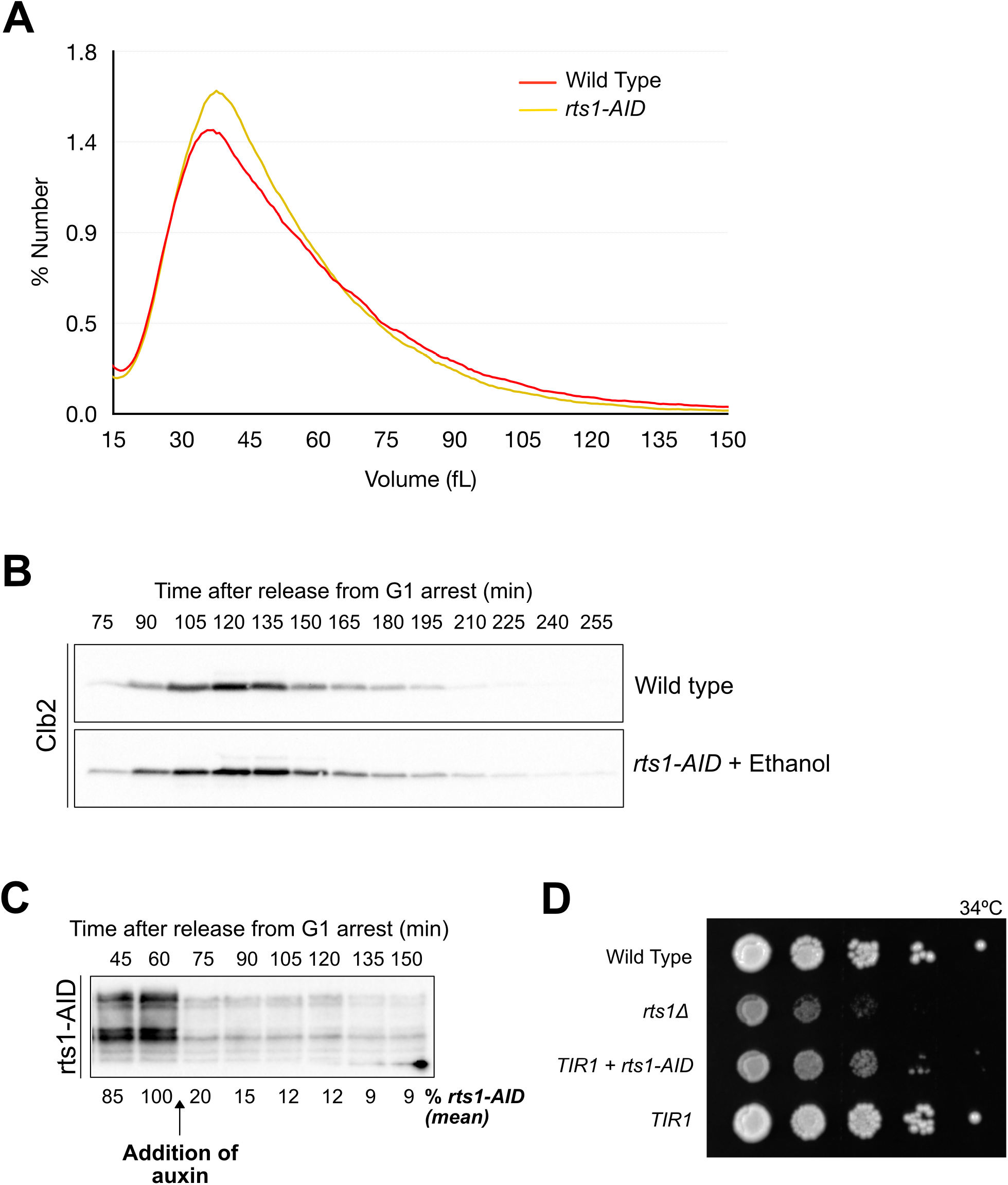
Characterization of *rts1-AID* cells. (**A**) Wild type and *rts1-AID* cells were grown to log phase in YPD medium and cell size was measured with a Coulter counter. (**B**) Wild type and *rts1-AID* cells were released from a G1 arrest in YPGE and the timing of entry into mitosis and the duration of mitosis were determined by assaying levels of the mitotic cyclin Clb2 by western blot. (**C**) *rts1-AID* cells were released from a G1 arrest in YPD medium and auxin was added at 60 minutes. Destruction of rts1-AID was assayed by western blot. The average rts1-AID signal was measured using BioRad Imagelab for 3 biological replicates and the mean percent of rts1-AID protein remaining at each time point is listed below each lane. (**D**) The rate of proliferation of *rts1*Δ and *rts1-AID* cells was tested by spotting a series of 10-fold dilutions of each strain on YPD medium containing auxin at 34°C.

**Figure S2:**
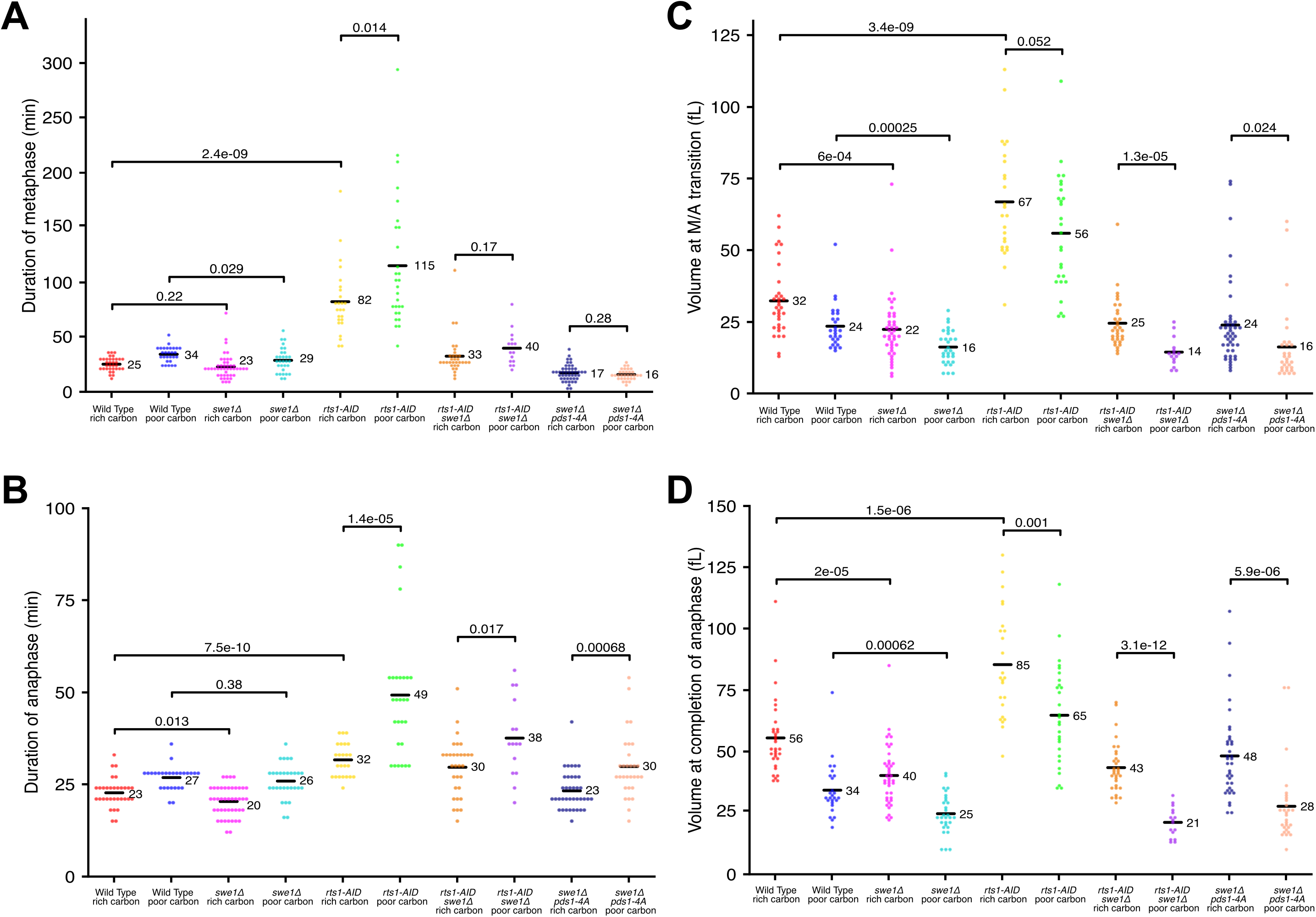
Dot plot versions of the data used to generate Figures 1,3, and 4. Black horizontal lines and adjacent numbers represent the average value for the data in each dot plot. The significance of the difference between two conditions is given as p-values above each plot.

**Figure S3:**
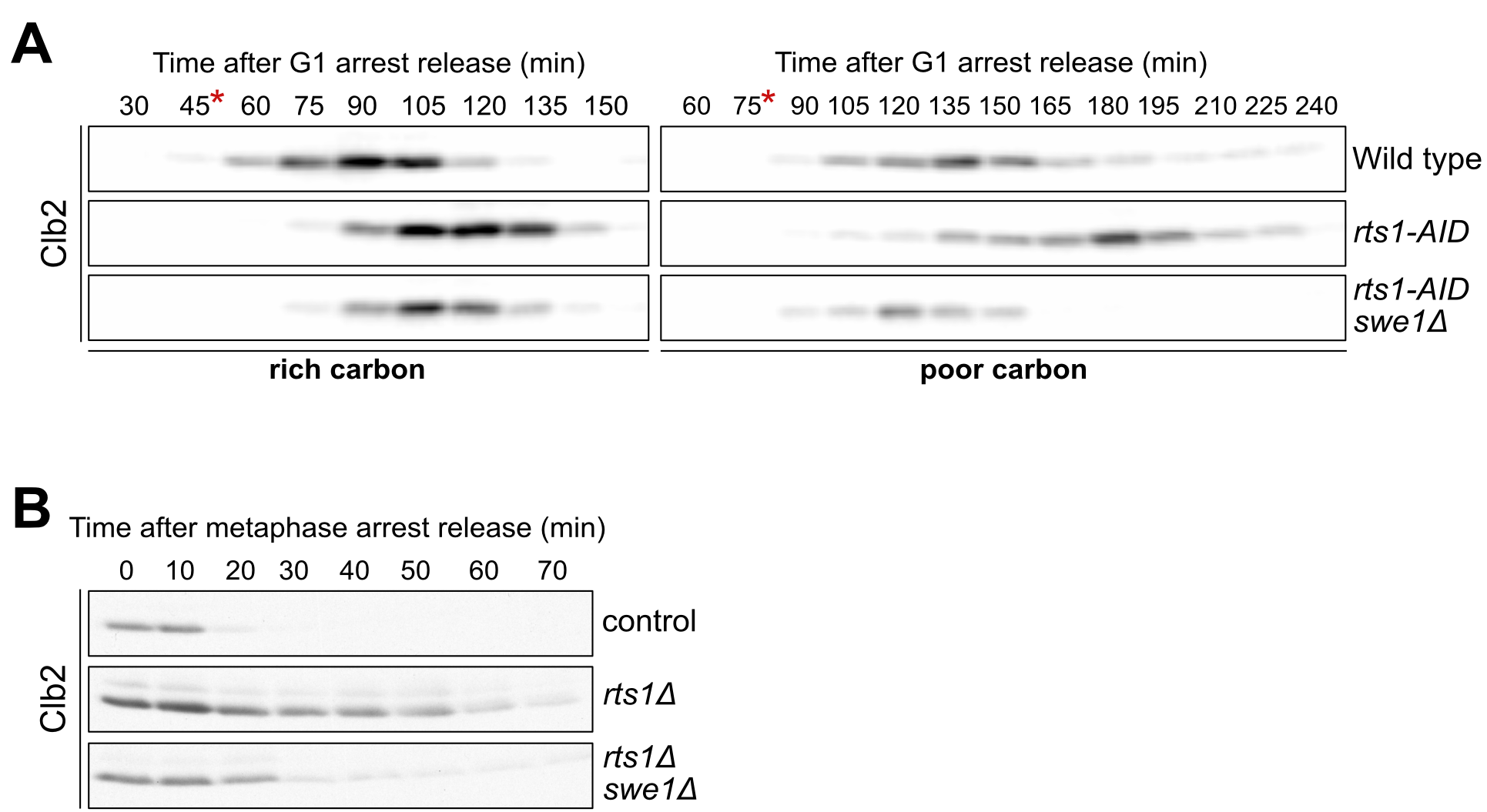
The increased duration of mitosis in *rts1-AID* cells is partially due to Cdk1 inhibitory phosphorylation. (**A**) Cells of the indicated genotypes, grown in YPD or YPGE, were released from a G1 arrest and auxin was added at 45 min (YPD) or 75min (YPGE). Addition of auxin is denoted by *. Levels of the mitotic cyclin Clb2 were assayed by western blot. (**B**) Cells of the indicated genotypes were arrested in metaphase by depletion of *GAL1-CDC20*. After release from the arrest, levels of the mitotic cyclin Clb2 were assayed by western blot.

